# Disordered Hippocampal Reactivations Predict Spatial Memory Deficits in a Mouse Model of Alzheimer’s Disease

**DOI:** 10.1101/2024.11.27.625618

**Authors:** S. Shipley, M.P. Abrate, R. Hayman, D. Chan, C. Barry

## Abstract

Alzheimer’s disease (AD) is characterised by progressive memory decline associated with hippocampal degeneration. However, the specific physiological mechanisms underlying hippocampal dysfunction in AD remain poorly understood and improved knowledge may aid both diagnosis and help identify new avenues for therapeutic intervention. We investigated how disruptions in hippocampal reactivations relate to place cell stability and spatial memory deficits in an AD mouse model. Using the APP knock-in mouse model ‘NL-G-F’, we conducted simultaneous behavioural and electrophysiological recordings in a radial arm maze. NL-G-F mice exhibited significant impairments in memory performance, demonstrated by an increased propensity to revisit arms, compared to wild-type controls. These memory deficits were associated with a reduction in the stability of hippocampal place cells, which occurred over short timescales and was accentuated across rest. While the rate of hippocampal reactivation events during rest was unchanged, the structure of these events was significantly degraded in NL-G-F mice. The decreased structure of reactivations was predictive of decreased stability in place cell firing. These findings suggest that disrupted reactivation sequences may act as a mechanism of memory disorder in AD.

## Introduction

The hippocampus plays a central role in memory for places and events, spatial navigation, and system-level memory consolidation (McClelland et al., 1995; Morris et al., 1982; Scoville & Milner, 1957; Squire, 1992). A key functional basis of the hippocampal role in navigation and spatial memory is the stable, spatially selective activity of place cells, primarily found in hippocampal regions CA3 and CA1 (Muller & Kubie, 1987; O’Keefe & Dostrovsky, 1971). The population activity of these cells is considered a core element of a cognitive map, providing a representation of self-location necessary to support flexible navigation to remembered goals (O’Keefe & Nadel, 1978). During quiescence, place cells engage in transient bursts of activity, reactivating temporally compressed trajectories through the environment (replay). These reactivation events, often synchronised with cortical activity (Ólafsdóttir & Barry, 2017; Rothschild et al., 2017), are widely considered a central mechanism supporting consolidation (Girardeau et al., 2009; Jadhav et al., 2012; Wilson & McNaughton, 1994). Within the hippocampus, replay events modulate synaptic efficacy, stabilising place fields (Buzsáki, 2015; O’Neill et al., 2008). Disruption of reactivations leads to place field destabilisation, underscoring the essential nature of these precise temporal sequences in maintaining spatial representations (Girardeau et al., 2009; Jadhav et al., 2012).

This understanding of the role of reactivation in memory consolidation has implications for understanding the nature of the memory deficit that is the hallmark symptom of early Alzheimer’s disease (AD) (Gauthier et al., 2021), which is in turn consistent with the hippocampus and associated medial temporal lobe regions being the first to manifest neurodegeneration and neuronal loss in AD (Braak & Braak, 1997; Braak & Del Tredici, 2015; Thal et al., 2002). The advent of anti-amyloid immunotherapies as the first wave of new therapies with the potential to modify disease and delay progression to dementia has increased the importance of early disease detection, given disease-modifying treatments are of greatest benefit in earliest stages of AD when the extent of irreversible neurodegeneration is lowest (Sperling et al., 2011). In this context, greater understanding of the physiological mechanisms underlying the earliest cognitive deficits in AD will benefit efforts to deliver tests with superior ability to detect the disease in its initial stages. However, identification of hippocampal pathophysiological changes would have benefits beyond early diagnosis alone, in that it would provide a mechanistic target against which to direct future symptom-targeted interventions.

Building upon our understanding of hippocampal function and its relevance to AD, studies in rodent AD models have reported a range of abnormalities in place cell firing characteristics. These include changes to spatial information, field size, and firing rate, which precede changes in other brain structures (Broussard et al., 2022; Cacucci et al., 2008; Cheng & Ji, 2013; Mably et al., 2017; Zhao et al., 2014). While there is a some variability in results arising from different mouse models, reflecting differences in the expression of AD molecular pathology and associated neuronal loss (Cheng & Ji, 2013; Ciupek et al., 2015; Jun et al., 2020) the most consistent finding across studies is the deterioration of place cell stability over various time-frames, both between and within recording sessions (Broussard et al., 2022; Ciupek et al., 2015; Mably et al., 2017; Zhao et al., 2014).

Given that hippocampal reactivation is critically involved in maintaining place field stability (Dupret et al., 2010; O’Neill et al., 2008), disrupted reactivation sequences may contribute to the reduction in field stability seen in AD. Earlier studies have identified a decrease in the quantity of sharp-wave ripples in several rodent AD models (Booth et al., 2016; Ciupek et al., 2015; Gillespie et al., 2016; Iaccarino et al., 2016; Jura et al., 2019; Witton et al., 2016). However, these transient features in the local field potential only indirectly measure reactivation events - they do not assess content or quality, and likely underestimate their occurrence. As such, the relationship between the actual content of reactivation - the information it conveys - and its impact on place cell stability remains unclear. Beyond this, the existing findings regarding reactivations require validation in modern knock-in AD models, which offer more physiologically relevant protein expression (Saito et al., 2014; Sasaguri et al., 2017). To better understand the link between hippocampal dysfunction and memory deficits in AD, three key aspects require further investigation: (1) the relationship between place cell instability and spatial memory impairments; (2) the temporal progression of these deficits throughout environmental exposure and experience; and (3) the direct mechanism and role of disrupted reactivations in mediating this relationship.

Here we investigated offline reactivations as a mechanism contributing to place field instability in an AD mouse model, and, in turn, the link to memory impairment. We explored place cell stability across different timescales in a familiar 8-arm radial maze while simultaneously evaluating two distinct forms of memory performance on the maze. To ensure clinical relevance, we employed an APP knock-in model, circumventing the problems associated with APP overexpression seen in many amyloid models of AD (Hsiao et al., 1996; Saito et al., 2014). In order to directly assess hippocampal reactivations, we identified epochs of multi-unit activity (MUA) from single cell recordings in CA1 during rest. We found an initial deficit in place cell stability in AD mice within a familiar environment, which improved over time. This instability was closely linked to impaired memory performance on the radial maze, particularly in the inability to recall previously visited arms within trials. Contrary to previous reports of sharp-wave ripples (Ciupek et al., 2015; Gillespie et al., 2016; Jura et al., 2019), analysis of reactivations during rest periods between behavioural sessions uncovered that there was no reduction in the overall number of reactivations. Rather, we observed a degradation in the structure of reactivations, which predicted the reduction in place cell stability. These findings illuminate the relationship between place cell activity, reactivation patterns, and memory performance. Our findings build upon previous studies by providing a complete link from cognitive deficits in AD, specifically memory impairments, with hippocampal place cell instability and disrupted sequences within reactivation events. This suggests that alterations in reactivation patterns could serve as an early indicator of hippocampal dysfunction and a potential target for therapeutic interventions in AD.

## Results

### NL-G-F mice show a reduced preference for unvisited arms in a radial arm maze task

Eight male APP^NL-G-F^/Chat-cre mice (from here on referred to as ‘NL-G-F’) and nine age matched wild-type littermate controls (two-sample t-test: M^WT^=12.22 months, Range^WT^=8,17; M^AD^=11.50 months, Range^AD^=7,15; t=0.50, p=0.63) were trained to run on an 8-arm radial arm maze until they were consistently running to the ends to receive rewards. During training, all arms were rewarded. Concurrently, extracellular electrodes (64 channels per mouse) were gradually lowered until they reached the CA1 cell layer (neural recordings performed in 15 of the mice; 8 wild-type and 7 NL-G-F). Once competency had been achieved and the cell layer reached, mice progressed to the experimental recordings (Fig.1A, Methods). Therefore, for all mice at the start of testing the maze was a familiar environment but the task was novel.

**Figure 1.**
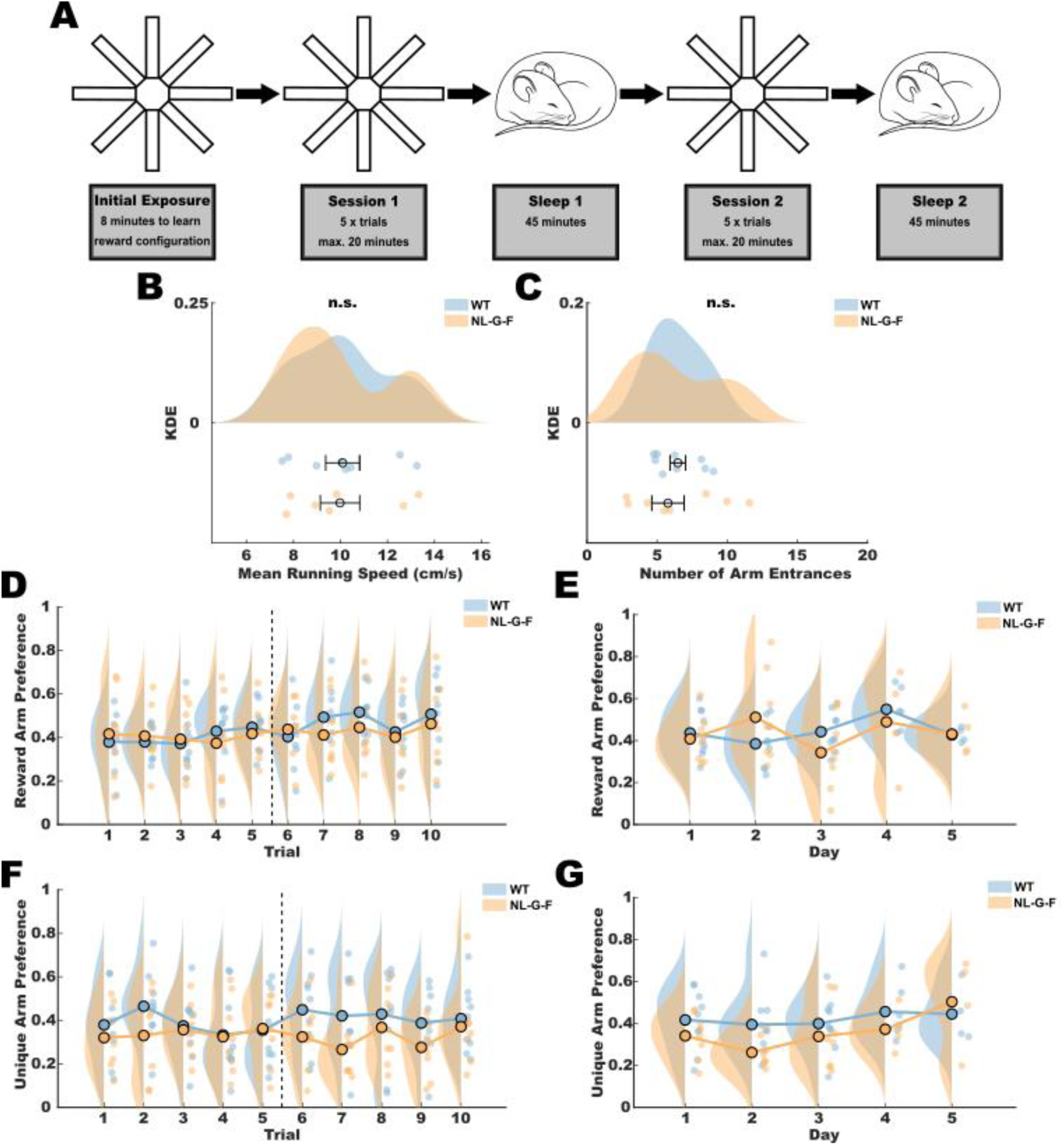
NL-G-F mice have a reduced preference for unvisited arms in an 8-arm radial arm maze task. **A.** Daily experimental procedure on the radial arm maze, which consisted of 8 arms (80cm x10cm) connected by an octagon (10cm sides). **B.** Mean running speed and **C.** number of arm entrances per trial was not different for WT and NL-G-F mice. Bars show mean and standard error, shaded area shows Kernel Density Estimate (KDE). n.s.= not significant in two-sample t-test. **D-E.** Mean and KDE of reward arm preference scores across trials **(D)** and days **(E)**. There was an overall increase in reward arm preference across trials (p<0.01) but no other significant factors. Dashed line represents the position of the sleep recording separating sessions 1 and 2. **F-G.** Same as **D-E** but for unique arm preference score. NL-G-F mice showed a decreased preference for visiting unique arms (p<0.05), and the preference decreased across trials (p<0.05) and days (p<0.01), though positive interaction effects existed between trial and day (p<0.01), and day and genotype (p<0.05). **D-G** significance tested using LME models (*Performance Score ∼ 1 + Trial*Day + Trial*Genotype + Day*Genotype + Trial:Day:Genotype + (1 | Mouse)).* **B,C,D,F.** *n^WT^*=9 mice, *n^NL-G-F^*=8 mice. **E,G.** Days 1-3: *n^WT^*=9, *n^NL-G-F^*=8. Day 4: *n^WT^*=6, *n^NL-G-F^*=4. Day 5: *n^WT^*=4, *_n_NL-G-F*_=4._

Mice completed a total of 3-5 test days (Supplementary Table 1). On each test day, four of the arms were randomly selected as reward arms. The reward configuration remained consistent throughout all 10 trials in a test day, but changed across days. Running behaviour was evaluated based on two parameters: mean running speed and the number of arm entries per trial. No significant differences were observed between NL-G-F mice and wild-type controls in either running speed (Fig.1B; two-sample t-test: M^WT^=10.09 cm/s, M^NL-G-F^=9.98 cm/s, t=0.10, p=0.92) or number of arm entries (Fig.1C; two-sample t-test: M^WT^=6.48, M^NL-G-F=^6.44, t=0.03, p =0.97).

To complete the task efficiently, mice should enter only the reward arms and avoid revisiting any arm, thus minimising travel distance and time. We assessed memory performance using two key metrics: the preference for visiting reward arms and the preference for visiting unique, previously unvisited, arms. For each of these, we developed a metric scored between 0 and 1, expressing animals’ preference for reward or unvisited arms whilst controlling for the total number of arm entrances (see Methods). The score represents the probability that an animal choosing the same number of arms at random would make a greater number of errors than observed. These scores allow us to quantify the animals’ ability to remember both the location of rewards and their previous choices, independent of their overall activity level. These metrics were analysed across 10 trials within each day, and across different days using linear mixed-effects models which represent the effect of a predictor variable on outcome, and accounted for individual variability among the mice .

Both wild-type and NL-G-F mice demonstrated an increase in their preference for reward arms over the course of the 10 trials within a day (Fig.1D-E, Supplementary Table 2; Estimate=0.026, SE=0.010, t(675)=2.667, p <0.01), indicating that they were capable of learning the reward configuration. However, there was no effect of genotype and interaction effects were not significant (Genotype: Estimate=0.188, SE=0.101, t(675)=1.869, p=0.062; Day: Estimate=0.034, SE=0.021, t(675)=1.636, p=0.102; Trial x Day: Estimate=-0.004, SE=0.003, t(675)=-1.078, p=0.281; Trial x Genotype: Estimate=-0.025, SE=0.014, t(675)=-1.729, p=0.084; Day x Genotype: Estimate=-0.052, SE=0.031, t(675)=-1.697, p=0.090).

In contrast to wild-type animals, NL-G-F mice were more likely to return to arms they had already visited (Fig.1F-G; Estimate=-0.243, SE=0.105, t(675)=-2.315, p=0.021). We also saw significant main effects of Trial and Day, with unique arm preference decreasing over time and trials (Trial: Estimate=-0.024, SE=0.010, t(675)=-2.413, p=0.016; Day: Estimate=-0.060, SE=0.021, t(675)=-2.892, p=0.004). Equally, significant interactions existed between trial and day (Estimate=0.01, SE=0.003, t(675)=3.02, p=0.002) and day and genotype (Estimate=0.070, SE=0.030, t(675)=2.296, p=0.022), but not trial and genotype (Estimate=0.017, SE=0.014, t(675)=1.177, p=0.240). Collectively, these results demonstrate that NL-G-F mice specifically failed to avoid revisiting arms. Although both genotypes showed an increased likelihood to revisit arms across trials and days, this was less pronounced in NL-G-F mice, as their score improved relative to wild-types.

### Deficits in place cell stability predict memory performance in NL-G-F mice

Place cells were recorded from mice navigating the radial arm maze (Fig.2A, Supplementary Fig.1; total across days: n^WT^=965, n^NL-G-F^=806), with key firing properties assessed only on the first test day to avoid repeated inclusion of cells (n^WT^=232, n^NL-G-F^=222). NL-G-F mice exhibited a reduced peak firing rate compared to controls (Fig.2B; two-sample t-test: M^WT^=10.11Hz, M^NL-G-F^=8.22Hz, t=3.66, p<0.001) and lower spatial information content of spikes (Fig.2C; two-sample t-test: M^WT^=1.21 bits/spike, M^NL-G-F^=0.80 bits/spike, t=5.42, p<0.0001), indicative of impaired spatial encoding. However, no significant difference in firing field size was observed between the genotypes (Fig.2D; two-sample t-test: M^WT^=40.11 bins, M^NL-G-F^=35.82 bins, t=1.63, p=0.103).

**Figure 2.**
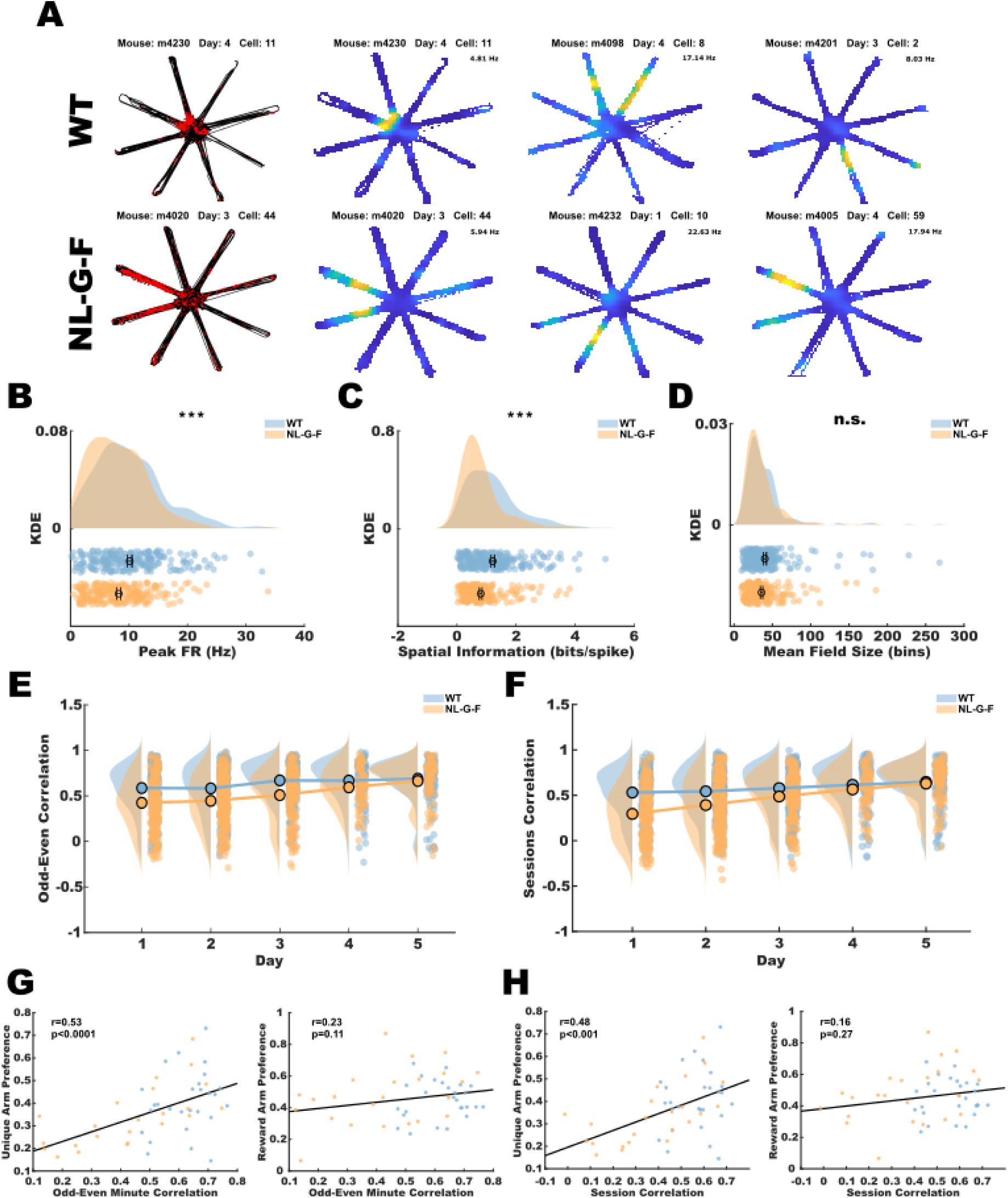
Reduced place field stability in NL-G-F mice corresponds to a decreased preference for unvisited arms. **A.** Spike raster plots (left) and ratemaps (right) of example place cells recorded in WT (top) and NL-G-F (bottom) mice. Rasters represent individual spike positions recorded over one day (concatenated across 10 trials). Ratemaps are colour coded from 0 to peak firing rate, labelled above each map. The first ratemap corresponds to the raster plot. **B.** Peak firing rate and **C.** Spatial Information of place cells were diminished in NL-G-F mice (t-test, p<0.001). **D.** Mean field area, averaged across all fields recorded from single cells, was not different between WT and NL-G-F mice (t-test, p=0.1). **B-D.** Cells recorded on the first test day across all mice (*n^WT^*=232, *n^NL-G-F^*=222). **E.** Odd-even minute rate map correlations were significantly lower in NL-G-F mice (p<0.001) and increased across days (p<0.01). **F.** Session-to-session (session 1 vs. 2) correlations of rate maps were also reduced in NL-G-F mice (p<0.0001) and showed an increase over days (p < 0.001), with NL-G-F mice improving more over time than WT mice (p<0.05). **E-F.** Pearson correlations of paired rate maps, excluding unvisited spatial bins. Statistical analysis was conducted with an LME model: *Correlation value ∼ 1 + Geno*Day + (1 | Mouse). n^WT^*=965 cell days, *n^NL-G-F^*=806 cell days. **G.** Place cell stability across odd and even minutes was significantly correlated with unique arm preference (left) but not reward arm preference (right). **H.** Place cell stability across sessions was also significantly correlated with unique arm preference (left) but not reward arm preference (right). **G-H.** Line represents spearman correlation. Each dot represents a recording day for a single mouse (*n*=50 days; colour coded by genotype for visualisation: *n^WT^=27*, *n^NL-G-F^=*23).

Place cell stability was evaluated using spatial correlations between ratemaps corresponding to two different time horizons: those composed of odd versus even minutes within a session, and those composed of the first versus second recording sessions of each day. Across session stability, reflecting longer time periods, was strongly reduced in NL-G-F mice (Fig.2F; Estimate=-0.28, SE=0.07, t(1765)=-4.25, p<0.0001), and improved across days (Estimate=0.02, SE=0.007, t(1765)=3.44, p<0.001). Notably, a significant interaction between genotype and day was observed, suggesting that while the stability of NL-G-F place fields was initially poor, it improved across test days (Estimate=0.02, SE=0.01, t(1765)=2.39, p<0.05).

Odd-even minute correlations were also lower in NL-G-F mice, reflecting reduced stability relative to wild-types over shorter time periods (Fig.2E; Estimate=-0.23, SE=0.06, t(1767)=-3.57, p<0.001). We also saw a primary effect of day, indicating that odd-even minute stability increased in both groups (Estimate=0.02, SE=0.007, t(1767)=3.12, p<0.01) - with no interaction between genotype and day (Estimate=0.007, SE=0.01, t(1767)=0.75, p=0.45).

Both stability measures showed a strong positive association with a preference for unique arms (Fig.2G,I, Supplementary Fig.2; Odd-Even Correlation: r(50)=0.55, p<0.0001; Session Correlation: r(50)=0.48, p<0.001) - indicating that animals with more stable place fields were less inclined to revisit arms. Conversely, no such relationship was observed with reward arm preference (Fig.2H,J; Odd-Even Correlation: r(50)=0.19, p<0.20; Session Correlation: r(50)=0.16, p=0.27). These findings highlight a specific link between the stability of hippocampal place cell representations and the ability to avoid revisiting arms within a given trial.

### Impaired Place Field Stability and Sleep-Dependent Consolidation in NL-G-F Mice

To investigate temporal variation in place cell stability between NL-G-F and wild-type mice, we generated separate minute-by-minute ratemaps and performed pairwise comparisons across all temporal delays within sessions. Correlations between ratemaps at different temporal delays were assessed through linear mixed-effects modelling (see Methods), allowing us to examine how stability changed over time and differed between genotypes.

Consistent with our previous analyses, overall, correlations were lower in NL-G-F mice (Fig.3A-B; NL-G-F Estimate=-0.098, SE=0.003, t(295980)=-3.24, p=0.001), corresponding to a general reduction in the place cell stability in NL-G-F animals. We also saw two additional main effects, a negative influence of temporal delay (Estimate=-0.005, SE=0.0003, t(295980)=-16.27, p<0.0001) and a positive effect of session (Estimate=0.04, SE=0.03, t(295980)=16.60, p<0.0001). Indicating, as expected, that place cell responses became increasingly dissimilar over more distant time points and that spatial responses tended to be more stable in the second session of the day.

**Figure 3.**
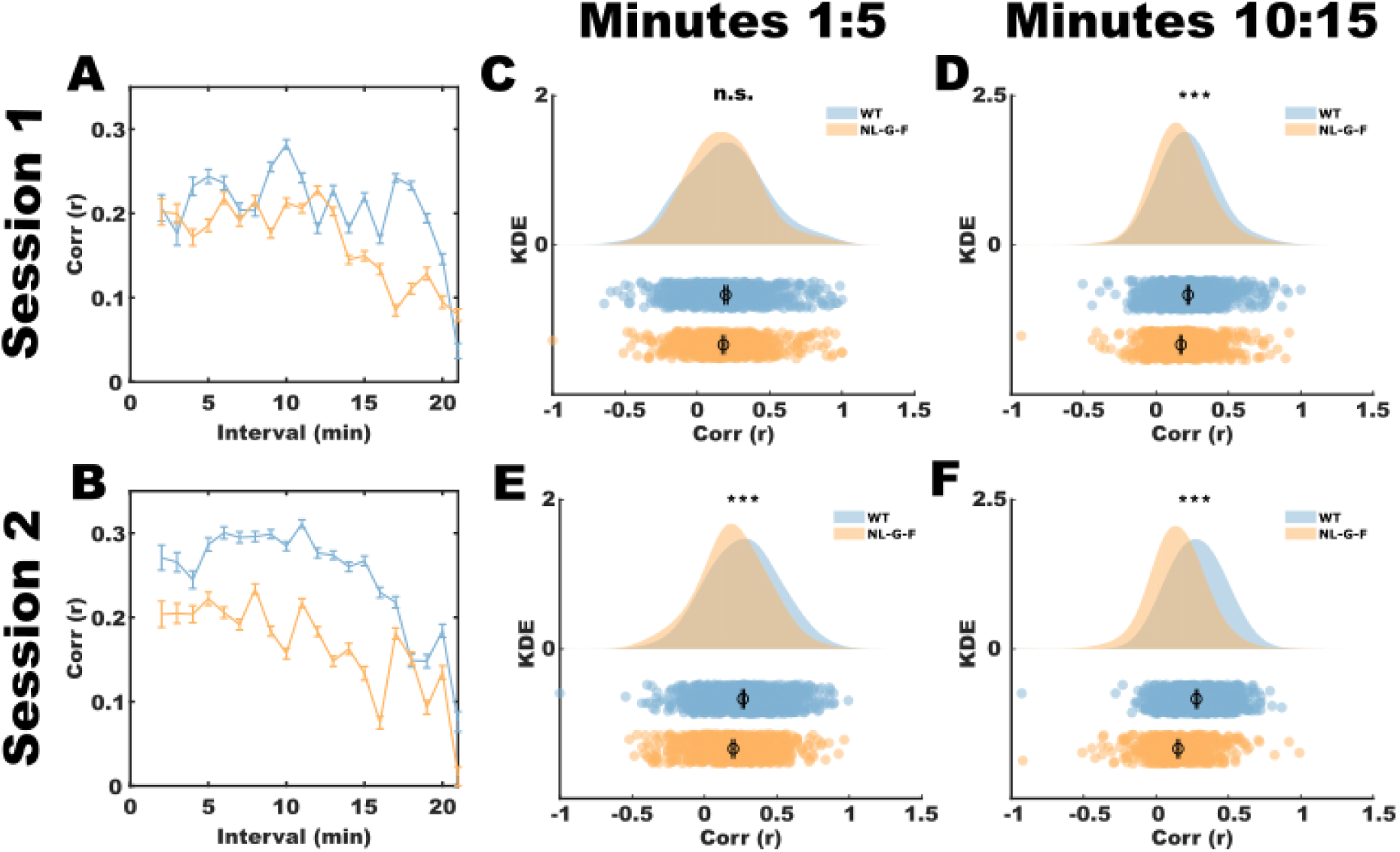
Place field stability degrades more rapidly and does not increase during rest in NL-G-F mice. **A,B.** Pearson correlations and SEM between individual one-minute ratemaps within session 1 **(A)** and 2 **(B)**. Both temporal delay between ratemaps and the NL-G-F genotype was predictive of decreased correlation (p<0.0001 and p=0.001 respectively). A significant interaction effect of genotype and session indicates an across session increase in stability in WT but not NL-G-F mice (p<0.0001). A small but significant interaction between temporal delay and genotype indicates a steeper decrease in WT mice (p<0.0001). LME model: *Correlation value ∼ 1 + Session*Genotype + Time Lag*Genotype + (1 | mouse)).* Ratemap correlations were performed on regions of overlapping paths, and excluded if there was no overlap (*n^WT^*=305688 , *n^NL-G-F^*=228781 one-minute ratemap pairs). **C.** At short temporal delays (1-5 minutes) in session 1, correlation values between WT and NL-G-F mice were not significantly different (*n^WT^*=690, *n^NL-G-F^*=663 ratemap pairs; p=0.27). **D.** At longer temporal delays (10-15 minutes) in session 1, NL-G-F mice exhibited decreased ratemap correlations compared to WT mice (*n^WT^*=947, *n^NL-G-F^*=740 ratemap pairs; p<0.0001). **E.** In session 2, following a rest session, NL-G-F mice had lower ratemap correlations at short temporal delays (*n^WT^*=949, *n^NL-G-F^*=791 ratemap pairs; p<0.01). **F.** Decreased ratemap correlations in session 2 were exacerbated over longer temporal delays (10-15 minutes; *n^WT^*=942, *n^NL-G-F^*=677 ratemap pairs; p<0.0001). **C-F.** p-values from 2-sample t-tests. Plots show individual Pearson correlation values and KDE. Bars show mean and SEM.

**Figure 4.**
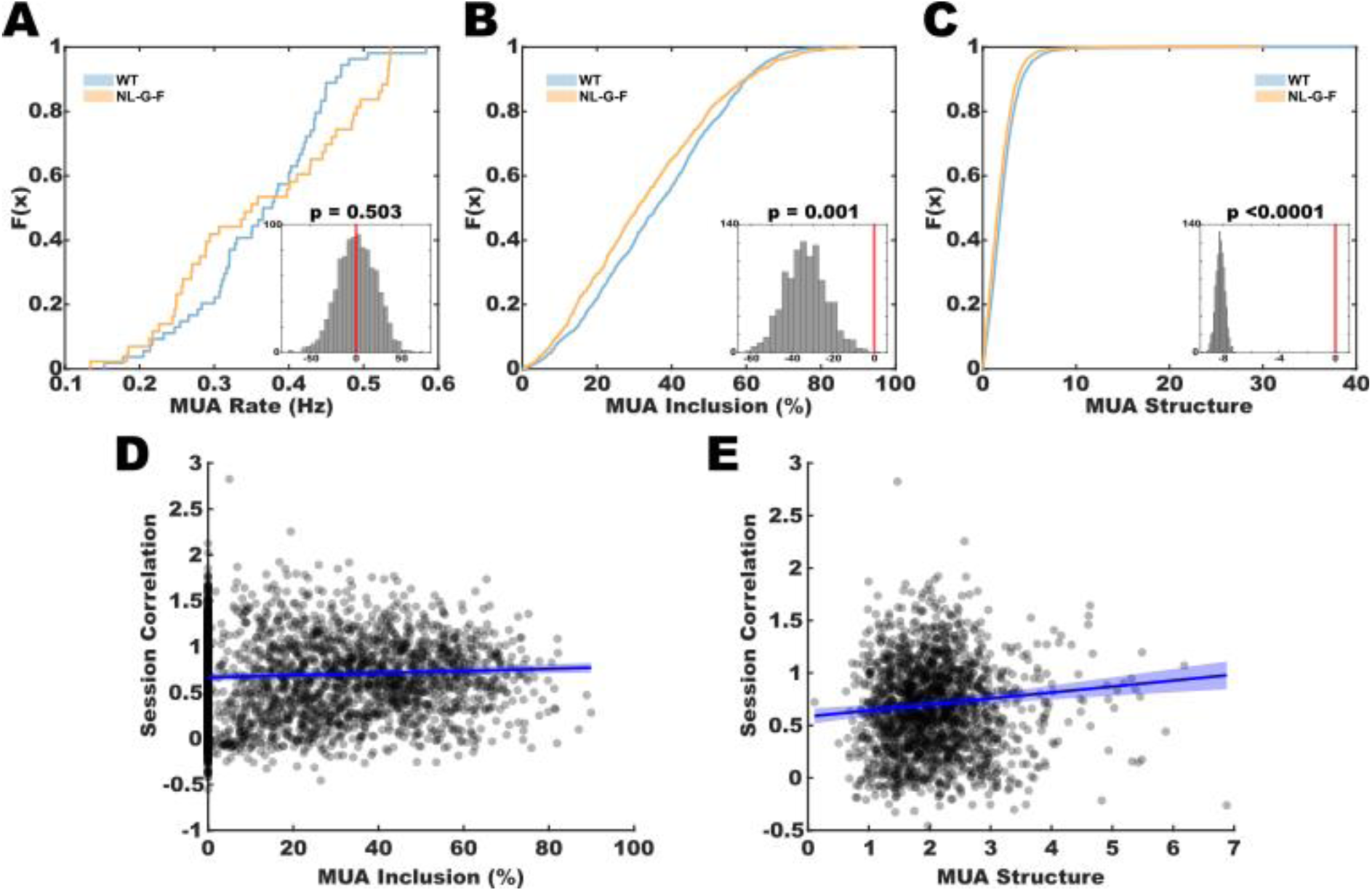
Cell recruitment in reactivations contributes to instability in place cell firing. **A.** Cumulative Distribution Frequency (CDF) plots of MUA rate in WT and NL-G-F mice. Difference in area under the curve (AUC) were tested for significance using a bootstrapping procedure (1000 iterations), with the resulting distribution (inset) being compared to zero (no difference; red line). There was no difference in MUA rate between genotypes (p=0.5). *n^WT^*=54, *n^NL-G-F^*=43 sleep recordings. **B.** Similar to A, for the percentage of MUAs to which each cell was recruited. MUA recruitment was reduced in NL-G-F mice (p=0.001). *n^WT^*=965, *n^NL-G-F^*=806 cell days. **C.** Similar to A,B. The level of structure within MUAs, measured as the likelihood of cell pair co-activity under a null hypothesis of statistical independence, was also reduced in NL-G-F mice (p<0.0001). *n^WT^*=48465, *n^NL-G-F^*=21921 cell pairs. **D-E.** Linear modelling demonstrated that recruitment to MUAs **(D)** and structure in MUAs **(E)** were both predictive of cell stability across sessions (z-transform of the Pearson correlation of ratemaps, p<0.0001). Plots show individual data points per cell (E: this is averaged across all relevant cell pair comparisons). *n^WT^*=965, *n^NL-G-F^*=806 cell days. Blue line represents the LME model estimate. LME model: *z-Scored Session Corr ∼ 1 + [MUA structure OR MUA inclusion] + (1 | Mouse)*.

Notably we found significant interactions between session and genotype (Estimate=-0.03, SE=0.004, t(295980)=-5.77, p<0.0001) and between temporal delay and genotype (Estimate=0.003, SE=0.0005, t(295980)=5.46, p<0.0001). These interactions suggest that the improvement in ratemap stability across sessions was less pronounced in NL-G-F mice compared to wild-type mice, while the decline in stability over increasing temporal delays was less steep in NL-G-F mice, although this effect was relatively small.

To further characterise these differences, we grouped correlations into 5-minute blocks and compared stability over short (<5 minutes; Fig.C,E) and long (10-15 minutes; Fig.D,F) temporal delays. During session 1, the stability of ratemaps separated by short temporal delays (<5 minutes) was not different between genotypes (M^WT^=0.20, M^NL-G-F^=0.18 r-value; two-sample t-test, t(1351)=1.11, p=0.27). However, ratemaps separated by 10 to 15 minutes were less stable in NL-G-F mice compared to wild-type mice (M^WT^=0.22, M^NL-G-F^=0.17 r-value; two-sample t-test, t(1685)=5.14, p<0.0001). This indicates that ratemaps in wild-type mice were stable over longer intervals. In session 2, following a sleep break, NL-G-F place fields were less stable than wild-type cells even at short temporal delays (<5 minutes; M^WT^=0.27, M^NL-G-F^=0.20 r-value; two-sample t-test, t(1738)=5.70, p<0.0001), with the size of the effect increasing over longer temporal delays (M^WT^=0.28, M^NL-G-F^=0.15 r-value; two-sample t-test, t(1617)=13.50, p<0.0001). These results suggest that, following rest, place fields from wild-type mice had higher initial levels of stability, and that stability was maintained over longer intervals.

Collectively, these findings indicate a general reduction in place cell stability across longer temporal lags within a session, an effect which was more prominent in NL-G-F mice. Furthermore, place fields in wild-type animals became more stable after the rest period between sessions, while NL-G-F mice failed to show this improvement. We therefore hypothesised that variations in sleep reactivations between genotypes might contribute to the observed disparities in ratemap stability.

### Coherent sleep reactivations stabilise the hippocampal map

The stabilisation of place fields has been attributed to reactivations of place cells during sleep (Dupret et al., 2010; O’Neill et al., 2008), suggesting a potential mechanism for the observed disparity between wild-type and NL-G-F mice. To explore this, we identified epochs of elevated multi-unit activity (MUA), which encapsulate reactivations (Ólafsdóttir et al., 2016, 2018), and assessed them for differences in quantity, cell recruitment, and structure.

The rate of MUA events was identical between wild-type and NL-G-F mice (MUA rate: M^WT=^0.36Hz, M^NL-G-F=^0.36Hz, bootstrapped differences in AUC: p=0.50). However, we observed a significant difference in how individual neurons participated in these events;place cells in NL-G-F mice were recruited to a lower proportion of events than in wild-type mice. (M^WT=^36.06%, M^NL-G-F^=33.06%, bootstrapped differences in AUC: p=0.001).

To assess how cell populations were recruited to MUAs, we developed an MUA structure score to evaluate the likelihood of cell pair co-activity under a null hypothesis of statistical independence. In essence, this quantifies the extent to which the observed joint probability of cell pairs being active in MUAs differed from the expected probability based on their individual activation frequencies (see Methods). A score of zero would indicate no difference from random combinations of cell pairs within MUAs, while higher scores suggest more structured, non-random co-activations. MUA structure in wild-type animals was significantly greater than in NL-G-F mice (M^WT^=2.24, M^AD=^1.91, bootstrapped differences in AUC: p<0.0001), suggesting that MUAs recorded from wild-type mice are more ordered, plausibly conveying more information.

The relationships between cell recruitment to MUAs, MUA structure, and place field stability between sessions were assessed using linear mixed-effects models (see Methods). These models demonstrated that both factors were significant positive predictors of across-session place field stability (MUA Structure: Estimate=0.06, SE=0.01, t(1725)=4.13, p<0.0001; MUA Inclusion: Estimate=0.003, SE=0.0005, t(1725)=5.00, p<0.0001). Together, these findings indicate that the ability to recruit and coordinate the firing of cell pairs within reactivations during rest plays a crucial role in maintaining stable neural activity across sessions. Thus, the degraded ability to recruit and structure cells into MUAs in the NL-G-F model of AD reveals a potential mechanism underlying the deterioration of place field stability and associated memory impairment.

## Discussion

In this study, we employed a multifaceted approach combining behavioural assessment in a radial arm maze task with simultaneous single-cell extracellular recordings to investigate hippocampal function, including reactivations, in a NL-G-F mouse model of Alzheimer’s disease (AD). We identified a direct relationship between place field stability and memory performance, both of which were impaired in NL-G-F animals. Notably, we observed a time-dependent deterioration of place field stability. Over short time frames, place fields in NL-G-F mice were stable comparable to those in wild-type animals. However, over longer periods, the stability of place fields in NL-G-F mice was markedly reduced relative to controls. This deficit was particularly pronounced following rest periods, after which the stability of place fields in wild-type animals increased whereas in NL-G-F mice it did not, resulting in a widening disparity between the two groups over longer temporal delays. To investigate the mechanisms underlying these differences, we examined reactivations during rest periods. We found that place cells were recruited to fewer multi-unit activity (MUA) events in NL-G-F mice, and that those MUAs were less structured, with cells reactivating in less consistent combinations. Linear modelling revealed that both MUA recruitment and MUA structure were predictors of across-session stability, suggesting a crucial role for coordinated reactivations in maintaining place field stability.

Understanding the neural mechanisms underlying memory impairment has major implications for AD management, not just in terms of yielding outcomes that may aid early diagnosis or track response to treatment but also in potentially identifying a target for future interventions aiming to restore normal physiological function. As such, a key aim of this study was to investigate the link between neural changes and memory performance. Using a radial arm maze task, we were able to evaluate different facets of memory, revealing a specific deficit in the preference for unique arms, rather than reward arms. This finding aligns with previous research that has shown reduced performance in alternating Y-mazes (Saito et al., 2014) and Morris-water-maze tasks (Mehla et al., 2019) in the same NL-G-F mouse model. Tasks such as these target short-term spatial memory, and have been shown to be more sensitive to impairment in AD than tests of reference memory (Stewart et al., 2011).

While the neural distinctions between these types of memory remain unclear, our findings suggest that the observed deficits are linked to the varying timescales required for learning different aspects of the task. In this study, preference for unique arms is an intra-trial measure of memory, relying on rapid, one-shot learning of ‘just visited’ but familiar locations. In contrast, preference for reward arms involves repeated learning across multiple trials within a day, showing improvement with practice. Consistent with this distinction, we saw that the stability of spatial representations reduced rapidly in NL-G-F mice, over short enough time periods to impact performance within a trial - such as odd vs even minutes. Indeed this reduction in place cell representation stability was explicitly correlated with a decreased preference for visiting unique arms. Conversely, across longer time periods - repeated days of exposure - the stability of place representations in NL-G-F animals recovered towards normal levels. Broadly this suggests that AD animals retain sufficiently normal neural dynamics to benefit from cumulative learning across multiple exposures to an environment, which may explain the relative preservation of reference memory observed by us and other researchers in these animals (Stewart et al., 2011).

Notably, the most pronounced difference between the two genotypes was in the stability of place cells after the 45-minute rest period - place cells in wild-type animals increased in stability, whereas those in NL-G-F mice did not. Hippocampal replay - compressed reactivations of hippocampal sequences (Louie & Wilson, 2001; Nádasdy et al., 1999; Wilson & McNaughton, 1994) - typically occur during quiescence and are known to play a central role in maintaining place field stability (Dupret et al., 2010; Jadhav et al., 2012; Karlsson & Frank, 2009; O’Neill et al., 2008). Prior work in AD rodent models has seen deficits in the occurrence of sharp-wave ripples - transient LFP phenomena that often co-occur with reactivations - in APP transgenic mouse models of AD (Iaccarino et al., 2016; Jura et al., 2019), tau models (Booth et al., 2016; Ciupek et al., 2015; Witton et al., 2016) and an APOE4 knock-in model (Gillespie et al., 2016). However this is far from a consistent finding across studies, likely because accurate measurement is confounded by differences in the oscillatory frequency of sharp-wave ripples between AD and wild-type animals (Benthem et al., 2020; Caccavano et al., 2020; Cayzac et al., 2015; Nicole et al., 2016).

Multi-unit activity (MUAs), the method used here, provides a more reliable indicator of reactivations than sharp-wave ripples used in previous studies, being a direct measure of hippocampal pyramidal cell activity (Ólafsdóttir et al., 2016; Widloski & Foster, 2024). Importantly, single unit recordings enabled us to directly assess the content of reactivations. Using this measure, we found no difference in the rate at which MUAs occurred between AD and wild-type animals. However, we observed that both the ability to recruit place cells into MUAs and the structure of cell co-firing within MUAs was reduced in NL-G-F animals. ‘This suggests that MUAs in NL-G-F mice conveyed less information than those in wild-type animals - factors which predicted place cell stability between sessions, which in turn was correlated with animals’ ability to avoid previously visited arms.

Whilst lacking a causal relationship, this triad of results strongly suggests the ability to recruit appropriate cells to MUAs is crucial for stabilising spatial maps during rest, which in turn provides a basis for subsequent short term spatial memory performance. The functional deficit in this AD model lies not in the ability of hippocampal networks to generate reactivations but in their precise ordering and internal structure. It follows that this aberrant activity is insufficient to engage the normal plastic processes required to stabilise the activity of place cells.

Further work is needed to fully clarify the causes of disrupted MUA structure in AD mice. A complete demonstration of this pathway, from mechanism to behaviour, would provide valuable insights into the memory impairments seen in AD. Moreover, it could reveal potential targets for symptom-specific therapeutic interventions, opening new avenues for understanding and treating dementia.

## Acknowledgments

We acknowledge funding from the following sources: SS from DeepMind Research Award, Cambridge Trust, Masonic Charitable Foundation; CB from Wellcome SRF, BBSRC; MPB from LIDo BBSRC; RH from Wellcome PRF; DC from NIHR, UK Dementia Research Institute, Wellcome, ARIA, Alzheimer’s Research UK.

## Author contributions

Author contributions: SS, CB and DC conceptualised the project design; SS, MPB conducted surgeries and collected data; SS and RH wrote code and analysed data; all authors wrote the paper.

## Methods

### Animals

All mice used in this study were from the APP NL-G-F/ChAT-cre genetic strain, generated by crossing the APP NL-G-F Alzheimer’s disease (AD) model (Saito et al., 2014) with ChAT-IRES-Cre (B6;129S6-Chattm2(cre)Lowl/J) mice. The ChAT-IRES-Cre line was established from a homozygous pair obtained from The Jackson Laboratory (ME, USA; Stock no: 006410). The colony was maintained by breeding pairs homozygous for ChAT-IRES-Cre and heterozygous for APPNL-G-F. APPNL-G-F homozygotes constituted the experimental group, while wild-type littermates served as controls. Electrophysiological recordings were performed on 15 male mice: 8 wild-type and 7 APP NL-G-F. An additional two mice, one APP NL-G-F and one wild-type, completed the behavioural task but were not implanted with electrodes.

Mice were aged to a minimum of 7 months prior to implantation (M^WT^=12.25 months, range^WT^=8,16; mean^-NL-G-F^=11.43, range^NL-G-F^=7,15; t_-(15)=_0.73, p=0.476). During this period, they were housed communally under a 12-hour reversed light-dark cycle. After surgery, each mouse was housed individually and food-restricted to 90% of their free-feeding body weight, or to 35g for those classified as obese. All procedures were carried out in accordance with institutional (University College London) and national ethical standards, following the UK Animals (Scientific Procedures) Act of 1986.

### Surgery

Mice were implanted with a single microdrive carrying 64 electrodes which were twisted into 16 separate tetrodes. Tetrodes were formed of twisted 12 μm polyimide coated nickel-chrome wire (California Fine Wire Co., CA, USA) which were platinum-plated to reduce impedances to below 150kΩ. Microdrives were formed of a single body (Axona Ltd., St Albans, UK) mounted to a screw which allowed all tetrodes to be advanced through the brain simultaneously.

Anaesthesia was induced in a chamber through which an isoflurane-oxygen mix (3ml/min) was administered. Once induced, analgesia was administered subcutaneously (carprieve – 5mg/kg, 1:9 solution in sterile saline). Mice were head-fixed on a stereotaxic frame and the anaesthetic gas mix was lowered to between 0.5 and 3 ml/min which was maintained throughout the surgery. Mice were placed on a heat-pad to maintain body temperature and were visually monitored for colour or breathing rate changes. Following surgery, Metacam (Meloxicam - 1mg/kg) was given in a Jelly suspension for a minimum of three days.

Following induction and placement in a stereotaxic frame, an incision was made along the midline to expose the skull, followed by the creation of three craniotomies. One craniotomy was used for the insertion of a screw attached to a metal pin which would be used for grounding, and the other two were used for the insertion of screws to stabilise the implant. A trephine drill was then used to create a craniotomy above the left hippocampus, where a single drive was implanted (coordinates: 2.0 mm posterior to bregma, 1.8 mm lateral to the midline, 0.8mm dorsal from the brain surface). Following insertion, a metal sheath was lowered over the exposed tetrodes for protection and the surgical site was covered with dental cement to fix the microdrive to the skull.

### Place Cell Screening

Following a one week post-surgery recovery period, mice were handled and habituated to the experimental room. The drive was then connected to the recording system and mice were allowed to forage freely in an open-field environment (80cm diameter). Neural and position data were recorded for 20-40 minutes, until the mouse had sufficiently covered the environment. During this time the neural data were reviewed for signatures of hippocampal activity (theta when running, sharp-wave ripples, pyramidal cell waveforms and bursts of cell activity). Spikes were initially clustered into cells using the automatic clustering algorithm KlustaKwik (Kadir et al., 2014) and reviewed manually in Tint (Axona Ltd., St Albans, UK) or using the Waveform software (d1manson.github.io/waveform). Over-clustering was corrected by merging clusters with similar waveforms and cross-correlograms indicating common spikes. Following this process, the number of cells with spatial fields, determined by eye, was assessed. If sufficient spatial cells were identified (a minimum of 20 spatial cells active in the open field environment), or if the number of spatial cells began to consistently decrease across recordings, the animal progressed into the experiment. If spatial cells were not identified, the tetrodes were lowered by turning the screw in the microdrive. Tetrodes were initially advanced up to 65µm between screening sessions, and later when neural patterns typical of the hippocampus were identified, they were advanced 30µm between screening sessions. This was performed up to three times in a day, with delays of at least two hours between each session.

### Behavioural Training

Experiments were performed on a radial eight-arm maze; arms were 80 cm x 10 cm, connected by an octagonal centre platform with sides of 10 cm and diameter of 24.7 cm. At the end of each arm was a reward well with raised edges such that a reward could not be seen from a distance. A second reward well was placed behind the end block of each arm which were all filled, to obscure odour cues.

Following a one-week post-surgery recovery period, mice were habituated to the experimental set up. On the first exposure mice were allowed to explore the radial arm maze without being connected to the recording tether. Subsequently, mice were connected and allowed to forage freely on the maze for eight-minute periods. Between exposures they were contained in a holding box for two minutes whilst the maze was cleaned and reward wells replenished. The holding box was made from a plastic flower-pot, with the bottom removed and split down one side so that the mouse could be contained and released without being picked up or untethered. The mouse was contained in this holding box in the centre of the maze, and the position of the side-opening was rotated randomly for each release to avoid route learning.

During the habituation phase a reward was provided on all arms of the maze to encourage exploration. If an animal visited an arm but did not eat the reward, the reward would be removed to discourage repeated visits. Habituation was performed on a minimum of two days and maximum of seven. The habituation phase was ended when the animal was able to consistently run to the ends of multiple arms within a single exposure and was regularly eating the reward. Once this habituation phase had been completed, mice progressed into the test phase.

### Behavioural Task

On each test day, reward wells on four of the eight possible arms contained a reward. The reward arms were selected randomly each day with the caveat that at least two arms must be different from the previous days configuration and there must be at least one unrewarded arm between the reward arms (i.e. not four consecutive arms). Once selected, this configuration remained for all trials within a day for that mouse. Reward arms were baited with half a chocolate vermicelli strand. Secondary reward wells behind all arm ends were also filled with a chocolate strand to mask the scent. On each day mice performed a total of ten trials, with the aim being to find all four reward arms on each trial.

At the start of a test day, the mouse was put in the holding box in the centre of the maze prior to being released for an 8 minute ‘exploration’ phase. This allowed the mouse to learn that day’s reward locations. They were then returned to the holding box for 2 minutes whilst the maze was cleaned and reward wells replenished. This re-setting procedure was performed between all exposures to the maze.

All test days consisted of two behavioural sessions and two sleep recordings. In all cases the animal progressed through the exploration phase, behavioural session one, sleep one, behavioural session two and sleep two on each day.

Each behavioural session consisted of five trials, each separated by the two-minute re-setting procedure. Each trial lasted for four minutes or until all four reward arms had been visited, whichever occurred first. At the end of each trial the mouse was returned to the holding box for two minutes and the maze re-set. During the behavioural task, the position of the mouse was tracked using an overhead camera and arm entrances were also manually recorded. Sleep recordings lasted 45 minutes. During this time, the mouse was held in a cylindrical rest box, placed on the centre platform of the maze to maintain position tracking. The mouse was monitored remotely during sleep recordings.

### Electrophysiology Recordings and Spike Sorting

Each microdrive contained two 32 channel Omnetics connectors to which Intan headstages could be attached. Headstages amplified and transmitted the neural signal to an Open Ephys acquisition board. Raw traces of the neural data were continuously reviewed in the Open Ephys GUI during recording, and were stored in the NWB format (Siegle et al., 2017).

To allow cells to be monitored across the two behavioural sessions as well as sleep recordings, neural data from the entire day’s recordings were combined. Initial clusters were assigned using the Kilosort 2.0 algorithm (Pachitariu et al., 2016; Stringer et al., 2019) which uses template matching to classify spikes into cluster based on putative cell identity. Clusters were then manually refined using Phy, an open-source spike sorting software package (Rossant et al., 2016). Amplitude, waveform and temporal autocorrelation of spikes were analysed to manually reassign spikes and to correct for over-clustering. After the spike sorting process was completed, spikes were reassigned to each trial in Matlab.

As the impact of Alzheimer’s disease on the firing properties of cells was unknown, a broad set of criteria was applied when allocating the cell assignments. Initially, clusters were categorised as either a pyramidal cell or interneuron based on the width of the waveform; a histogram of all peak to trough waveform durations was calculated and two clear peaks identified. The lower limit for inclusion of a cell as pyramidal was set to the minima between these two peaks (400µs). All cells with a waveform duration less than this threshold were categorised as interneurons.

The subset of cells considered putative pyramidal cells were assessed for spatial firing properties to determine whether they would be classified as a place cell. Firing fields were classified as areas on the maze of at least 9 connected 2cm bins, in which the firing rate of the cell exceeded twice its mean firing rate. In addition to having at least one firing field, cells were required to have a peak firing rate greater than 1Hz but less than 50Hz, and a mean firing rate of less than 5Hz to be considered a place cell (based on Tanni et al., 2022).

### Position Tracking

Headstages were fitted with two infra-red LEDs which provided a signal of the mouse’s location which was tracked using an overhead night-vision infrared webcam. An X,Y position was recorded using the PosTracker plugin for Open Ephys (<u>github.com/rhayman/PosTracker</u>) and inserted into the NWB data file. A position data point was stored at a rate of approximately 30Hz though there was some variability. To correct for this, position was interpolated and matched to the neural signal using a nearest-neighbour search of timestamps for the respective data sources.

Occasionally, the camera would not detect the LED for short periods of time or would detect an alternate light source and store that as the XY coordinate. This produced jumps in the position data. A post-hoc removal of any jumps greater than one standard deviation from the mean position change between timestamps was performed in Matlab and replaced through interpolation of neighbouring data points.

The rate of change in position was calculated using Pythagoras’ theorem and used to determine the running speed. Speed was calculated for each time stamp, adjusted for the pixels per meter recorded by the camera, and averaged to determine the distance travelled in centimetres per second.

### Ratemaps

Both position data and spikes were filtered for running speed so that only periods when the mouse was moving at a rate greater than 2 cm per second were included. Position samples were grouped into 2cm x 2cm bins and firing ratemaps were calculated by dividing the number of spikes in each spatial bin by the cumulative dwell time in that bin. Ratemaps were then smoothed with a Gaussian smoothing kernel (σ=4cm). The mean rate was calculated as the firing rate averaged across all spatial bins of the ratemap. The peak firing rate was the firing rate in the maximal spatial bin.

### Task Performance Scores

Behavioural performance was evaluated using scores based on unique arm preference and reward arm preference. These scores accounted for the number of choices made by a mouse during each trial by comparing the observed data to a shuffled distribution of error probabilities. Separate shuffled distributions were generated for both unique arm preference and reward arm preference scores. Each score was calculated as the inverse of the summed probability of making the observed number of errors or fewer, given the number of arm entrances.

Statistical analysis of behaviour was performed using linear mixed effects modelling, to account for individual variations between mice and assess the relative contributions of multiple variables. For both performance scores (unique arm preference and reward arm preference) the following model was used:
*(Performance Score ~ 1 + Trial*Day + Trial*Genotype + Day*Genotype + Trial:Day:Genotype + (1 | Mouse))*

### Spatial Information

Spatial information values were calculated for each cell, and provided a measure of the spatial specificity of that cell’s activity. Cells that fire broadly have a low spatial information value when compared to cells that have highly condensed areas of activity in the environment (firing fields). To avoid any impact of novelty, the spatial information was calculated using the neural data recorded during the second session on each test day.

As is standard, the spatial ratemaps used to measure spatial information were calculated using the adaptive binning method laid out in Skaggs *et al*. (1996); the firing rate in each bin was calculated by expanding a circle around the bin until the conditions of the following equation were met:

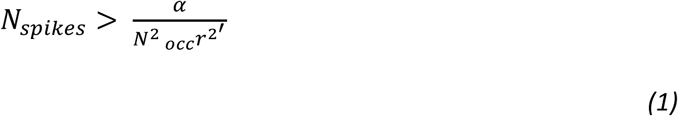

Where *N_occ_* is the number of occupancy samples falling within the circle, *N_spikes_* is the number of spikes occurring within the circle, *r* is the radius of the circle in bins and *α* is a scaling parameter, set at 200 in this instance. The value within each bin is therefore *Rs · N_spikes_ / N_occ_* where *Rs* refers to the position sampling rate.

Spatial information values were then attained using the method described by Skaggs *et al*. (1992). The information in bits per second was calculated using the formula:

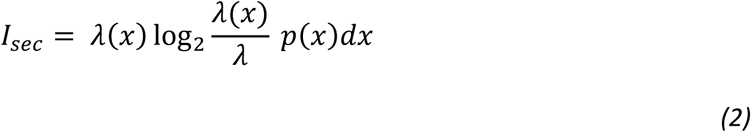

Where *x* is the spatial location and *p*(*x*) is the probability density of the mouse being in that location. λ(*x*) is the mean firing rate at position *x*. The overall mean firing rate of the cell is calculated as:

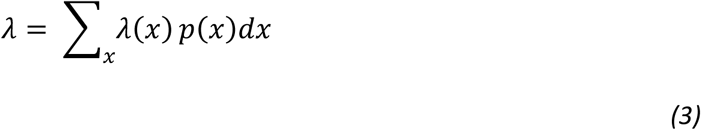

The spatial information in bits per spike can then be calculated for each cell using the equation:

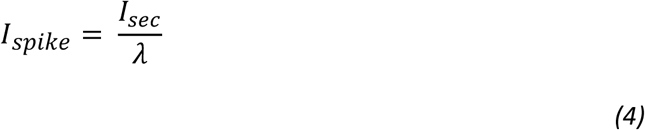

### Spatial Correlations

To assess place cell activity stability, we used spatial correlations to directly compare rate maps (R. Muller & Kubie, 1989). For across session stability, separate rate maps were generated for sessions one and two on each recording day for each place cell. These were compared using bin-wise Pearson correlations, excluding unvisited bins. Odd-even stability was assessed by generating separate rate maps for data recorded in ’odd minutes’ (e.g., minutes 1, 3) and ’even minutes’ (e.g., minutes 0, 2), then correlating these as before. To assess temporal dynamics of place cell stability, individual rate maps were created for every minute of recording across both sessions. Within a single session, all pairwise comparisons at different time lags were performed using the same correlation method.

Stability change across days, trials and between genotypes was assessed using linear mixed effects modelling. For both across session correlations and odd-even correlations, the following model was used:
LME model = *Correlation Value ~ 1 + Genotype*Date + (1 | Mouse)*

Linear mixed effects modelling was also used to assess the temporal dynamics of stability change within sessions, using the following equation:
LME model = *Correlation Value ~ 1 + Session*Genotype + Time Lag*Genotype + (1 | Mouse))*

### MUA Detection

Reactivation events were identified as epochs in which the multi-unit activity (MUA) of all place cells was increased, reaching three standard deviations above the mean activity within the sleep recording. To assess the changing level of multi-unit activity across the recording, a histogram of the spike activity of all cells, with 10ms time bins, was generated and smoothed with a Gaussian kernel (σ=15ms). The activity level was z-scored and potential MUA events were identified as periods of activity which exceeded 3 standard deviations above the mean.

Peaks of activity passing this threshold were identified, and the boundaries of the MUA were extended in both directions until the firing rate returned to baseline, providing the outer limits of the MUA. The total time between these limits was used to measure the duration of the MUA event. For inclusion as an MUA event, the duration was required to be greater than 40ms. For a cell to be considered active in an MUA, it must have fired at least one action potential within the duration of the MUA. These criteria were selected in line with previous studies (Bush et al., 2022; Ólafsdóttir et al., 2016, 2018).

### MUA Structure

To determine the structure of MUA content, we developed a score measuring the extent to which place cell co-firing differed from chance expectations assuming statistical independence. We defined the probability of the *i^th^* cell being active in any given MUA as:

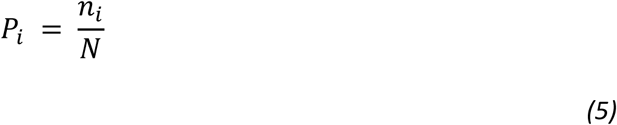

Where *n_i_* is the number of MUA events cell i is active in and N is the total number of MUA events in that sleep session.

The expected probability that any given pair of cells (*i* & *j*) - would be active in the same MUA events, assuming statistical independence, is then the joint probability:

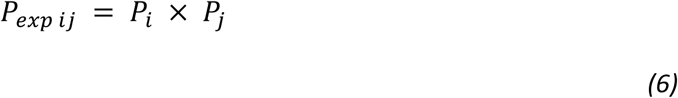

We determined the observed probability of cells *i* and *j* co-activating within MUAs as:

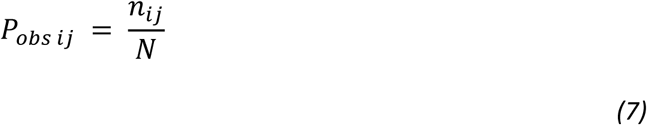

Where *n_ij_* is the number of MUA events in which both *i* and *j* were active. To quantify the departure of observed data from expected probabilities, we calculated a z-score and take the absolute value:

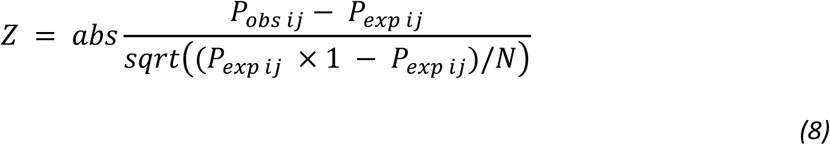

Where *N* is the total number of MUA events. Higher absolute Z-values indicate greater structure in MUA content, with 0 suggesting no structure (observed matches expected).

The relationship between MUA structure and the z-transform of the session correlation was assessed using the linear mixed effects model:
LME model = *z-Scored Session Correlation ~ 1 + MUA structure + (1 | Mouse)*

