## Supplementary Material for "Disordered Hippocampal Reactivations Predict Spatial Memory Deficits in a Mouse Model of Alzheimer’s Disease"

**WT**

| Mouse | Day | Age | Place Cells |
|---|---|---|---|
| m0168 | 1 | 12 | - |
| m0168 | 2 | 12 | - |
| m0168 | 3 | 12 | - |
| m0168 | 4 | 12 | - |
| m0168 | 5 | 12 | - |
| m4098 | 1 | 16 | 65 |
| m4098 | 2 | 16 | 52 |
| m4098 | 3 | 16 | 51 |
| m4098 | 4 | 16 | 41 |
| m4098 | 5 | 16 | 57 |
| m4101 | 1 | 16 | 47 |
| m4101 | 2 | 16 | 47 |
| m4101 | 3 | 16 | 44 |
| m4101 | 4 | 16 | - |
| m4101 | 5 | 16 | - |
| m4201 | 1 | 8 | 39 |
| m4201 | 2 | 8 | 58 |
| m4201 | 3 | 8 | 43 |
| m4201 | 4 | 8 | - |
| m4230 | 1 | 12 | 23 |
| m4230 | 2 | 12 | 21 |
| m4230 | 3 | 12 | 40 |
| m4230 | 4 | 12 | 58 |
| m4230 | 5 | 12 | 83 |
| m4376 | 1 | 17 | 20 |
| m4376 | 2 | 17 | 31 |
| m4376 | 3 | 17 | 32 |
| m4376 | 4 | 17 | 14 |
| m4578 | 1 | 11 | 14 |
| m4578 | 2 | 11 | 23 |
| m4578 | 3 | 11 | 15 |
| m4604 | 1 | 9 | 10 |
| m4604 | 2 | 9 | 9 |
| m4604 | 3 | 9 | 14 |
| m4605 | 1 | 9 | 14 |
| m4605 | 2 | 9 | - |
| m4605 | 3 | 9 | - |

**NL-G-F**

| Mouse | Day | Age | Place Cells |
|---|---|---|---|
| m0166 | 1 | 12 | - |
| m0166 | 2 | 12 | - |
| m0166 | 3 | 12 | - |
| m0166 | 4 | 12 | - |
| m0166 | 5 | 12 | - |
| m4005 | 1 | 15 | 20 |
| m4005 | 2 | 15 | 23 |
| m4005 | 3 | 15 | 27 |
| m4005 | 4 | 15 | 24 |
| m4005 | 5 | 15 | 30 |
| m4020 | 1 | 14 | 59 |
| m4020 | 2 | 14 | 61 |
| m4020 | 3 | 14 | 52 |
| m4020 | 4 | 14 | 51 |
| m4020 | 5 | 14 | 54 |
| m4202 | 1 | 7 | 9 |
| m4202 | 2 | 7 | 19 |
| m4202 | 3 | 7 | 15 |
| m4232 | 1 | 12 | 51 |
| m4232 | 2 | 12 | 56 |
| m4232 | 3 | 12 | 60 |
| m4602 | 1 | 10 | 7 |
| m4602 | 2 | 10 | 18 |
| m4602 | 3 | 10 | - |
| m4602 | 4 | 10 | - |
| m4602 | 5 | 10 | - |
| m4609 | 1 | 11 | 17 |
| m4609 | 2 | 11 | 15 |
| m4609 | 3 | 11 | 25 |
| m4610 | 1 | 11 | 59 |
| m4610 | 2 | 11 | 54 |
| m4610 | 3 | 11 | - |

**Supplementary Table 1. Number of place cells recorded across mice and days.** Mouse ID, days into the experiment, age (months) and number of place cells for each experimental day. Days where no place cell number is recorded were included in behavioural analyses only.

| Equation | Figure | Estimates | p Value |
|---|---|---|---|
| *Reward Arm Performance Score ~ 1 + Trial*Day + Trial*Genotype + Day*Genotype + Trial:Day:Genotype + (1 \| Mouse))* | 1D-E | Trial: 0.026 | **<0.01** |
| | | Genotype: 0.188 | 0.062 |
| | | Day: 0.034 | 0.102 |
| | | Trial x Day: -0.004 | 0.281 |
| | | Trial x Genotype: -0.025 | 0.084 |
| | | Day x Genotype: -0.052 | 0.09 |
| *Unique Arm Performance Score ~ 1 + Trial*Day + Trial*Genotype + Day*Genotype + Trial:Day:Genotype + (1 \| Mouse))* | 1F-G | Trial: -0.024 | **<0.05** |
| | | Genotype: -0.243 | **<0.05** |
| | | Day: -0.060 | **<0.01** |
| | | Trial x Day: 0.01 | **<0.01** |
| | | Trial x Genotype: 0.017 | 0.24 |
| | | Day x Genotype: 0.070 | **<0.05** |
| *Odd-Even Correlation value ~ 1 + Geno*Day + (1 \| Mouse)* | 2E | Genotype: -0.23 | **<0.001** |
| | | Day: 0.02 | **<0.01** |
| | | Day x Genotype: 0.007 | 0.45 |
| *Across Session Correlation value ~ 1 + Geno*Day + (1 \| Mouse)* | 2F | Genotype: -0.28 | **<0.0001** |
| | | Day: 0.02 | **<0.001** |
| | | Day x Genotype: 0.02 | **<0.05** |
| *One-Min Ratemaps Correlation value ~ 1 + Session*Genotype + Time Lag*Genotype + (1 \| mouse))* | 3A-B | Genotype: -0.098 | **0.001** |
| | | Time Lag: -0.005 | **<0.0001** |
| | | Session: 0.04 | **<0.0001** |
| | | Session x Genotype: -0.03 | **<0.0001** |
| | | Time Lag x Genotype: 0.003 | **<0.0001** |
| *z-Scored Session Correlation ~ 1 + MUA inclusion + (1 \| Mouse)* | 4D | MUA Inclusion: 0.003 | **<0.0001** |
| *z-Scored Session Correlation ~ 1 + MUA Structure Score + (1 \| Mouse)* | 4E | MUA Structure: 0.06 | **<0.0001** |

**Supplementary Table 2. Linear mixed effects models used throughout the paper.** All 'Genotype' effects are given as differences in NL-G-F data from WT baseline. In all cases, individual variability across mice was controlled for.

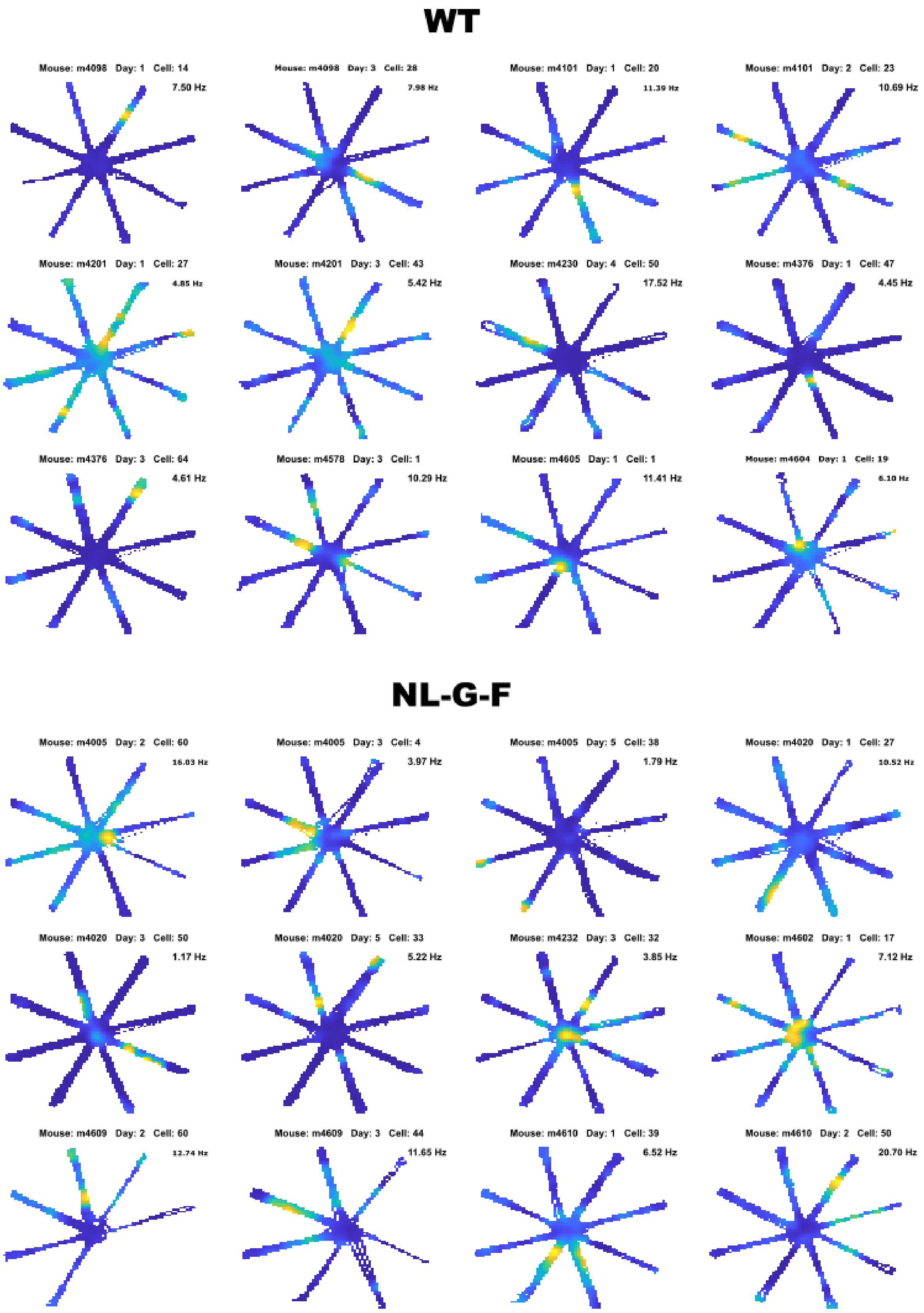


**Supplementary Figure 1. Example ratemaps of place cells recorded on the radial-arm maze.** Mouse ID, recording day and cell ID are given above each ratemap. Peak firing rate is provided at the top-right of each plot. Colours are scaled separately, from zero to the peak firing rate, for each cell. Unvisited bins are left white.

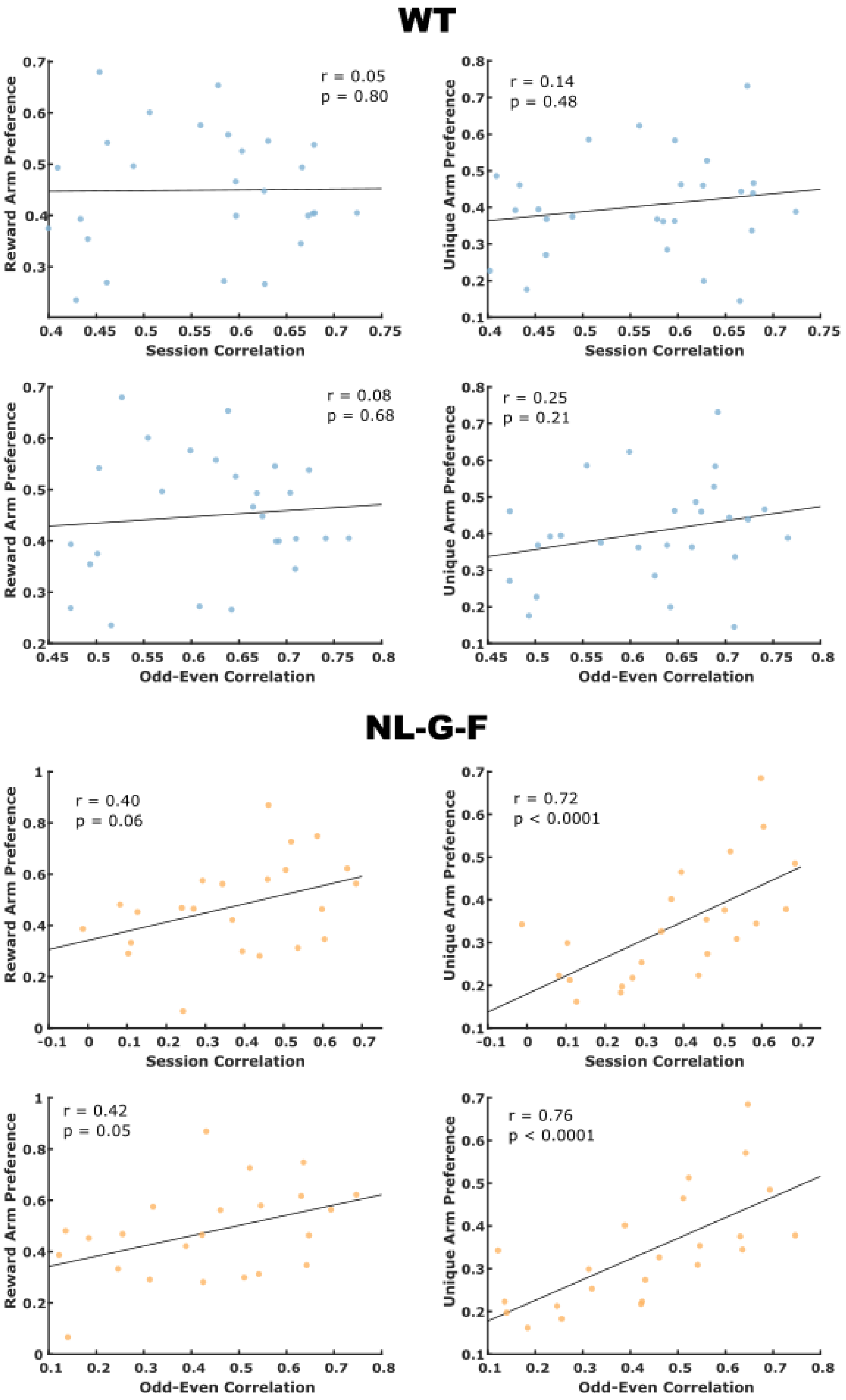


**Supplementary Figure 2. Correlation between stability measures and memory performance scores.** Significant positive correlations exist for NL-G-F mice between both measures of stability (Odd-Even Minute correlations and Across Session correlations) and unique arm preference scores. WT mice show less variability in stability scores, though a non-significant positive trend remains. Similar trends are seen for NL-G-F mice between stability and reward arm preference scores.  Line represents spearman correlation. Each dot represents a recording day for a single mouse
